# Non-invasive Free-breathing Gating-free Extracellular Cellular Volume Quantification for Repetitive Myocardial Fibrosis Evaluation in Rodents

**DOI:** 10.1101/2025.07.11.664411

**Authors:** Devin R. E. Cortes, Thomas Becker-Szurszewski, Sean Hartwick, Muhammad Wahab Amjad, Soheb Anwar Mohammed, Xucai Chen, John J. Pacella, Anthony G. Christodoulou, Yijen L. Wu

## Abstract

**Background:** Interstitial myocardial fibrosis is a crucial pathological feature of many cardiovascular disorders. Myocardial fibrosis resulting in extracellular volume (ECV) expansion can be non-invasively quantified by cardiac MRI (CMR) with T1 mapping before and after gadolinium (Gd) contrast agent administration. However, longitudinal repetitive ECV measurements are challenging in rodents due to the prolonged scan time with cardiac and respiratory gating that is required for conventional T1 mapping and the invasive nature of the rodent intravenous lines.

**Methods:** To address these challenges, the objective of this study is to establish a fast, free-breathing, and gating-free ECV procedure with a non-invasive subcutaneous catheter for in-scanner Gd administration that can allow longitudinal repetitive ECV evaluations in rodent models. This is achieved by (1) IntraGate sequence for free-breathing gating-free cardiac imaging, (2) non-invasive subcutaneous in-scanner Gd administration, and (3) fast T1 mapping with varied flip angle (VFA) in conjunction with (4) triple jugular vein blood T1 normalization. Additionally, full cine CMR (multi-slice short-axis, long-axis 2-chamber, and long-axis 4- chamber) was acquired during the waiting period for comprehensive systolic cardiac functional and strain analysis.

**Results:** We have successfully established a non-invasive fast ECV quantification protocol to enable longitudinal repetitive ECV quantifications in rodents. Non-invasive subcutaneous Gd bolus administration induced reasonable dynamic contrast enhancement (DCE) time course reaching a steady state in ∼ 20 min for stable T1 quantification. The free-breathing gating-free VFA T1 quantification scheme allows for rapid cardiac (∼2.5 min) and jugular vein (49 sec) T1 quantification with no motion artifacts. The triple jugular vein T1 acquisitions (1 pre-contrast and 2 post-contrast) immediately flanking the heart T1 acquisitions enable accurate myocardial ECV quantification. Our data demonstrated that left-ventricular myocardial ECV quantifications were highly reproducible with repeated scans, and the ECV values (0.25) are comparable to reported ranges among humans and rodents. This protocol was successfully applied to ischemic reperfusion injury model to detect myocardial fibrosis that was validated with histopathology.

**Conclusion:** We have established a simple, fast, non-invasive, and robust CMR protocol in rodents that can enable longitudinal repetitive ECV quantifications for cardiovascular disease progression. It can be used to monitor disease regression with interventions.

**Graphical Abstract:** 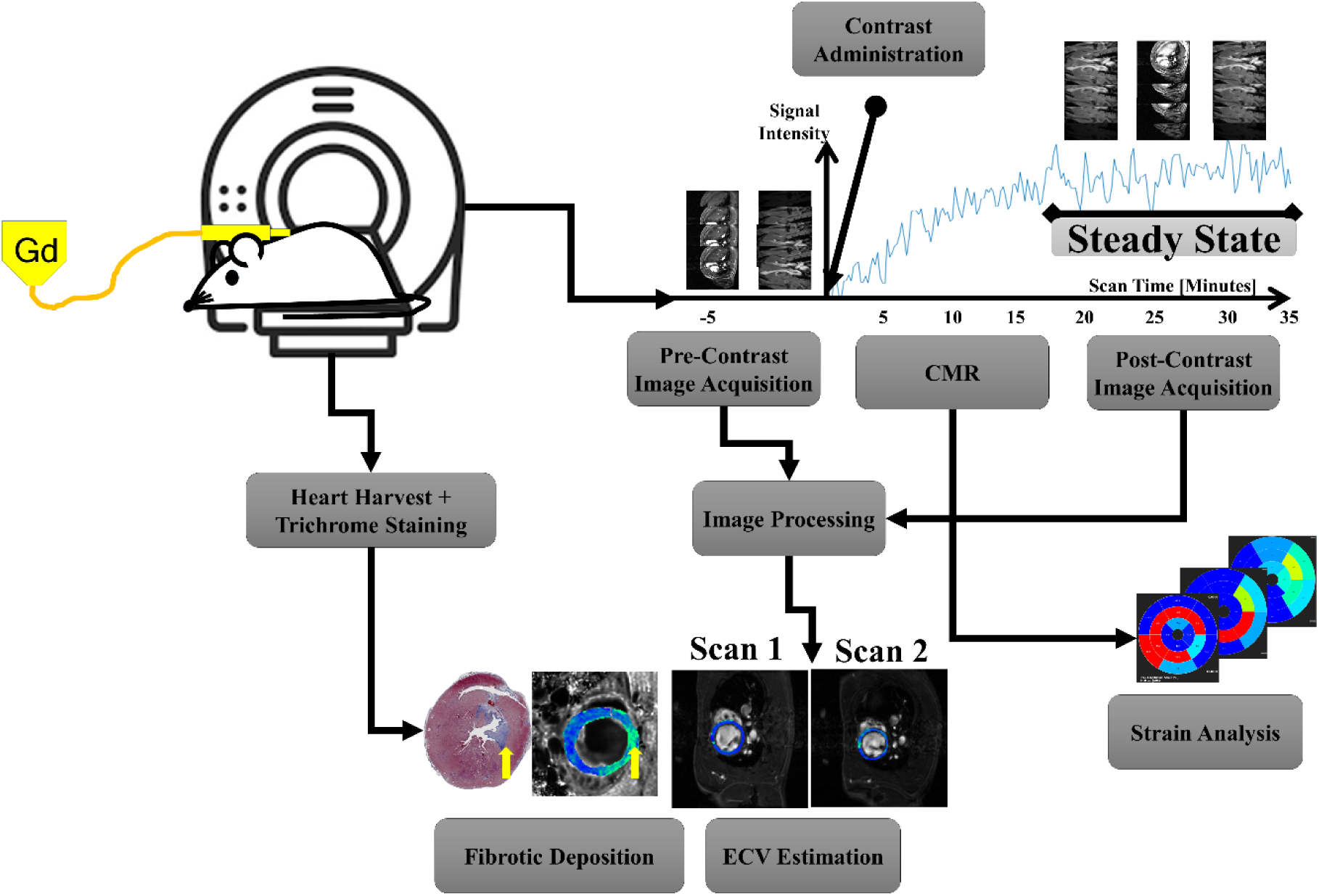

## 1. Introduction

Interstitial myocardial fibrosis [1–3] is a crucial hallmark of cardiovascular diseases, such as hypertrophic cardiomyopathy (HCM) [4–6], congenital heart disease (CHD) [7–9], and heart failure with preserved ejection fraction (HFpEF) [10–14]. Myocardial fibrosis resulting in extracellular volume (ECV) expansion can be non-invasively quantified by cardiac MRI (CMR) with T1 mapping before and after gadolinium (Gd) contrast agent administration [15–22]. As ECV expansion is progressive [14] and reversible, it is a therapeutic target and can be a surrogate biomarker for therapeutic trials [1, 23].

Animal models are indispensable in modeling human diseases for mechanistic understanding and testing therapeutic efficacy [24–26]. Rodent models are suitable for cardiovascular research [24, 27, 28] because rodents and humans share similar anatomy with 4- chamber hearts and similar great arteries and veins, comparable physiology, and conserved biological processes, genes, gene modifies, and molecular pathways [24, 29–31]. Compared to large animals, rodents have much shorter breeding cycles (18-21 days) and weaning ages (21 days), facilitating time efficient studies and cost reduction for animal housing. More importantly, the availability of inbred lines and transgenic strains makes mechanistic investigations of genetic and epigenetic regulations possible.

However, longitudinal repetitive ECV measurements are challenging in mice and rats. Common human T1 mapping methods with inversion recovery (IR) schemes [32–34], such as modified Look-Locker inversion recovery (MOLLI), shortened MOLLI (ShMOLLI), and saturation recovery single-shot acquisition (SASHA), usually require breath-holding and electrocardiogram (ECG) gating to eliminate cardiac and respiratory motion artifacts. Rodent CMR with respiratory and ECG gating can prolong the scan time under anesthesia, which can compromise physiological and hemodynamic functions. Currently available MRI-compatible rodent ECG systems are not robustly shielded from magnetic field distortion. ECG waveforms often get distorted in the magnet, making accurate and effective ECG gating problematic.

Additionally, in order to quantify myocardial T1 with and without Gd, it requires an intravenous (i.v.) line to administer Gd inside the magnet. Repetitive i.v. lines are challenging for rodents.

Tail or jugular vein catheters are not MRI compatible and can be easily clotted or collapsed. Femoral i.v. lines are invasive and terminal, thus not feasible for longitudinal repetitive injections.

To address these challenges, the objective of this study is to establish a free-breathing gating-free ECV procedure with non-invasive subcutaneous catheter for in-magnet Gd administration that can allow longitudinal repetitive ECV evaluations in rodent models. We leveraged the IntraGate sequence with varied flip angle (VFA) for fast quantitative T1 mapping without needing ECG nor respiratory gating [35, 36]. Additionally, left ventricular (LV) blood is commonly used to normalize myocardial T1 values along with hematocrits to quantify ECV [15–21]. However, VFA-measured-T1 values in blood are altered by turbulent flow in-slice and especially through-slice flow that introduces proton spins into the LV excitation region prior to the steady state. We have successfully overcome this issue and established a robust and reproducible ECV protocol in rodents with triple jugular vein T1 mapping using a non-invasive subcutaneous Gd administration.

## 2. Methods

### 2.1 Animals

Eight male Sprague Dawley (CD® IGS, Strain Code 001) rats were purchased from Charles River Laboratories. Animals were housed and cared for according to the animal protocol (IACUC protocol # 25056602) approved by the Institutional Animal Care and Use Committee, University of Pittsburgh. Animals were housed in cages with centralized filtered air/clean water supply system in a secure AALAS certified vivarium in the Rangos Research Center of the Children’s Hospital of Pittsburgh of UPMC. Animal care is provided seven days a week based on the NIH Guide for *the Guide for the Care and Use of Laboratory Animals.* Animals were kept in a 12:12 hour dark/light cycle with *ad libitum* food and water.

All data was generated and analyzed in accordance with ARRIVE [37] guidelines, including protocols for blinding, randomization, counterbalancing, inclusion of proper controls, and appropriate statistical power.

To test reproducibility of the method, each rat was repetitively imaged on 2 different days.

### 2.2 Anesthesia for animal imaging

All rats received general inhalation anesthesia with Isofluorane for *in vivo* imaging. A rat was placed into a clear plexiglass anesthesia induction box that allowed unimpeded visual monitoring of the animals. Induction was achieved by administration of 2-3% isofluorane mixed with oxygen for a few minutes. Depth of anesthesia was monitored by toe reflex (extension of limbs, spine positioning) and respiration rates. Once the plane of anesthesia was established, the rat was maintained with 1-2 % isofluorane in oxygen via a designated nose cone then the rat was transferred to the designated animal bed for MRI.

The core body temperature was monitored by a rectal optical probe and was maintained by circulating warm water inside the animal bed or feedback-controlled MR-compatible air heater module for small animals (SA Instruments. Inc. Model 1025). Respiration is monitored with a pneumatic pillow sensor coupled to a pressure transducer.

### 2.3 Subcutaneous catheter for Gd administration

A subcutaneous (subQ) catheter (Mckesson IV Catheter 25G x 0.75”, Product Number: 854664) was placed on the backside of each rat (Fig.1A). The catheter was then connected to a micro bore extension set (Braun 36 in, Reference Number: V6203) to enable Gd administration while the animal was inside the MRI scanner.

**Figure 1.**
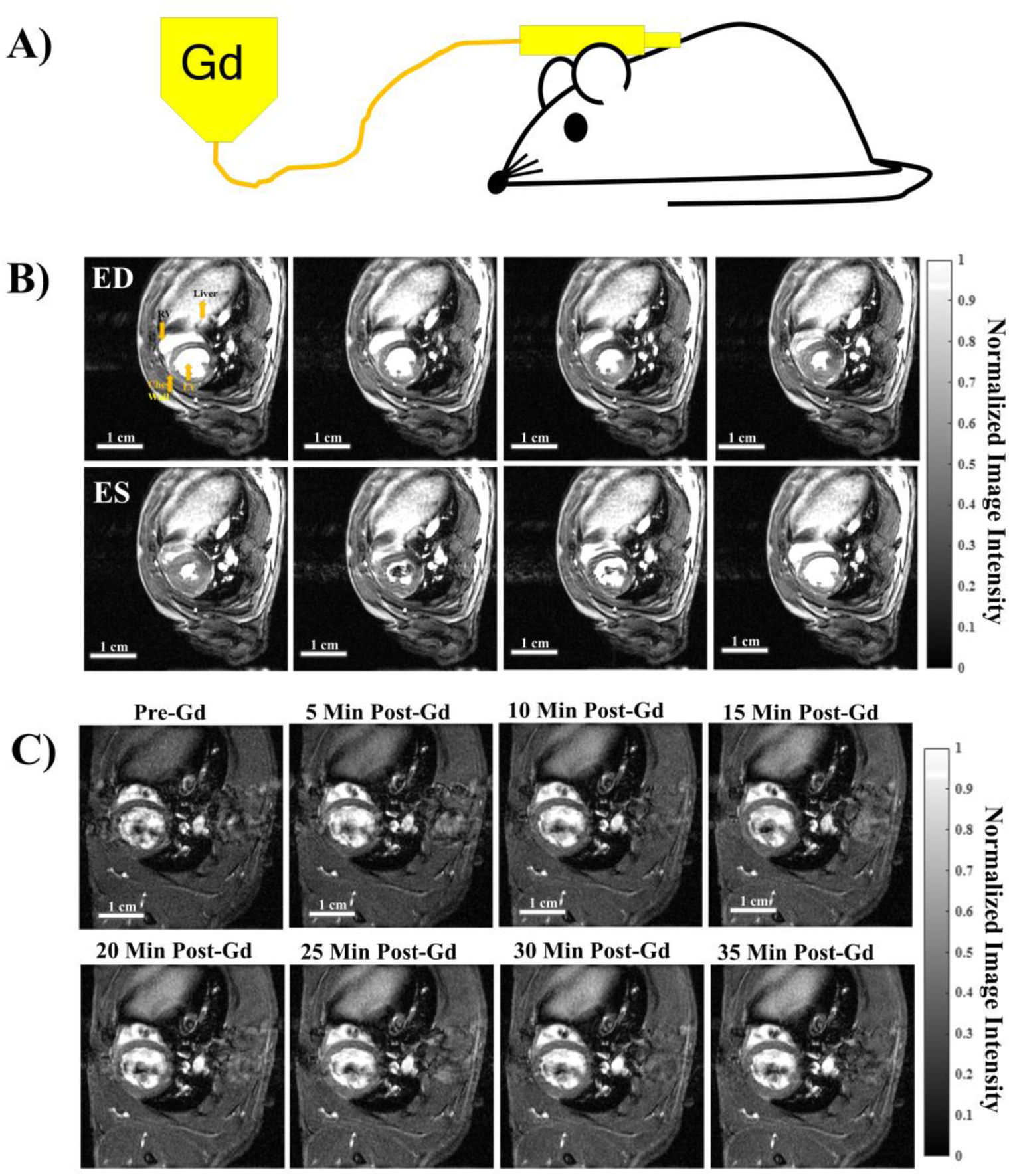
Free-breathing gating-free cine and dynamic contrast enhancement (DCE) time series with subQ Gd administration. (A) A single bolus of Gd was administered via a subcutaneous (subQ) catheter on the back of the animal inside the scanner to enable pre- and post-contrast acquisitions. (B) Cine time series of a full cardiac cycle acquired by IntraGate with FA = 28°, showing 8 out of the 20 cardiac phases. ED = end diastole. ES = end systole. (C) Dynamic contrast enhancement (DCE) time series at ED phase showing the overall increase in contrast from the time of Gd bolus injection.

A single bolus of clinical grade Gadobenate Dimehlumine (MultiHance®, Bracco Diagnostics, Inc. NDC 0270-5164-12) with a dosage of 0.2 mmol/kg bodyweight was injected via a subcutaneous catheter at time = 0 (Fig.2B).

**Figure 2.**
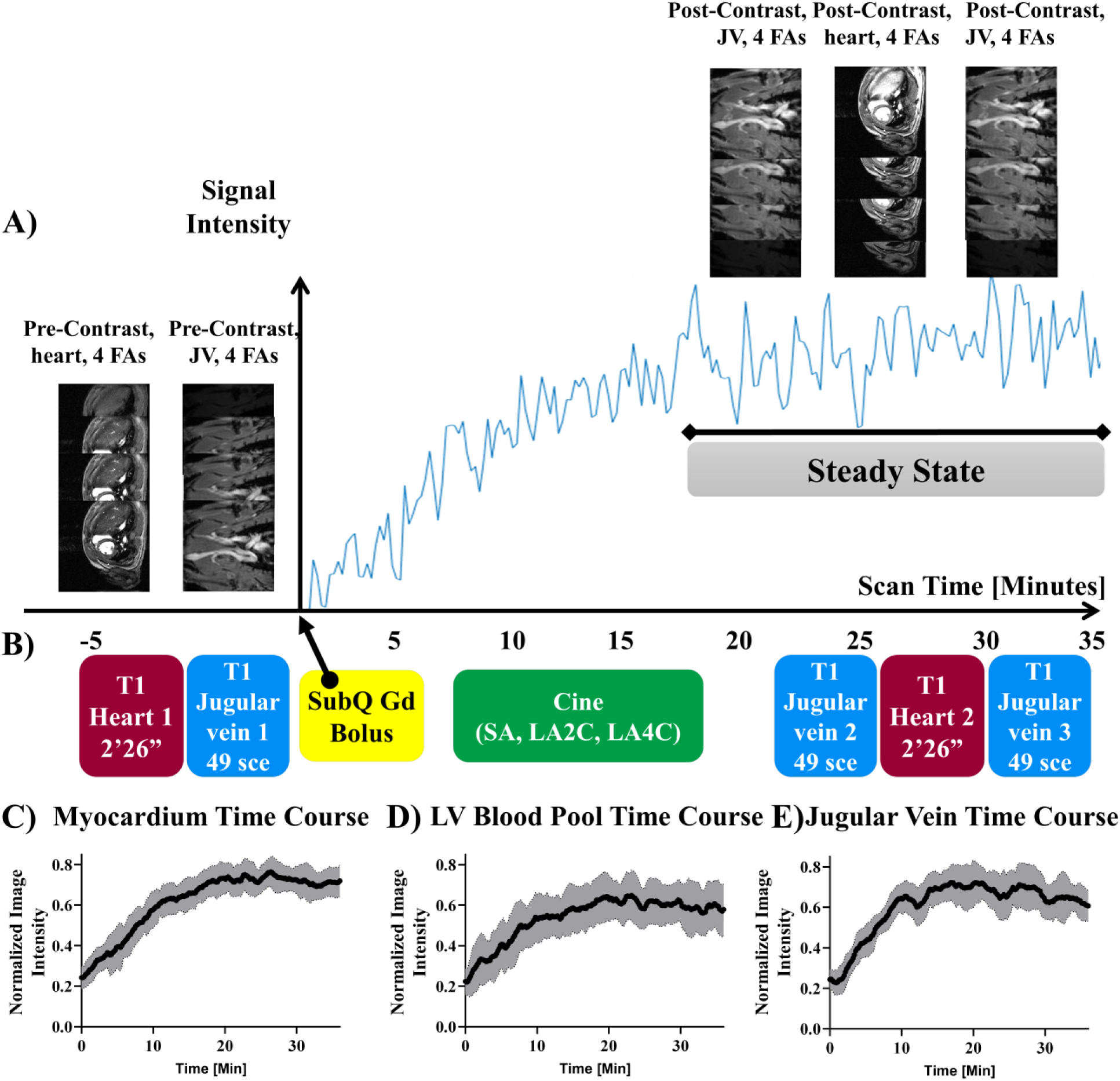
Image acquisition schema and DCE time course. (A) Temporal LV myocardial signal profile with dynamic contrast enhancement (DCE). Y-axis: signal intensity of LV myocardium. X-axis: time (minutes). The time point of the Gd bolus administration is set to 0. The LV myocardial signals reached steady state around ∼20 minutes post-Gd bolus. (B) Imaging acquisition scheme. Yellow blocks: Gd bolus of 0.2 mmol/Kg bodyweight. Red blocks: T1 mapping for the heart. Blue blocks: T1 mapping for jugular veins. Green block: cine CMR for multi-slice SA, LA2C, and LA4D. Prior to Gd injection, VFA heart and jugular vein image series were acquired with four different flip angles (FA): 3°, 19°, 22°, and 28°. After the Gd injection, it is critical to wait 20 minutes to acquire the next set of post-Gd image sets to allow for steady-state contrast. When the steady state contrast was achieved, next sets of images with 4 FAs were acquired in the following order: jugular vein, heart, and jugular vein again. Two sets of jugular vein images are averaged together for blood T1 quantification. (C-E) Time courses of the dynamic contrast time-series taken from (C) LV myocardium, (D) LV blood pool, and (E) jugular vein. All tissue and blood source reached the steady-state by 20 min. Y-axis is normalized image intensity. Black lines represent average signal intensity of all rats (n=8) and grey shaded regions represent standard deviation of each time point.

### 2.4 Hematocrit

A blood sample was collected via facial vein using a 5.5-mm sterile blood lancet (Goldenrod Animal Lancet). The blood was sealed with clay in a micro-hematocrit capillary tube and spun in a micro hematocrit centrifuge for 5 minutes to separate the blood cells and plasma.

Hematocrit was then measured as a ratio of the final plasma volume to the total blood volume.

### 2.5 CMR acquisition

All *in vivo* CMR was performed using a Bruker BioSpec 70/30 USR spectrometer (Bruker BioSpin MRI, Billerica, MA) operating at a 7-Tesla field strength with a 35-mm quadrature volume coil for both transmission and reception. Fast free-breathing-no-gating CMR was acquired with IntraGate sequence, a fast low-angle shot (FLASH)-based pulse sequence with retrospective gating without needing ECG or respiratory triggering for acquisition [35, 36]. Cardiac and respiratory motions were disentangled by the Bruker IntraGate interface for cardiac and respiratory cycles for retrospective imaging reconstruction.

*Dynamic contrast enhancement (DCE) time course*: T1-weighted DCE time course (Fig.1B) after a single bolus Gd injection at time = 0 was acquired with the following parameters: Field of view (FOV) = 5 × 5 cm, slice thickness (SLTH) = 1.5 mm, sampled with an acquisition matrix of 256 × 256, in-plane resolution of 0.195 mm, echo time (TE) = 3.06 msec, repetition time (TR) = 5.65 msec, flip angle (FA) = 10°, number of repetitions (NR) = 3000, and a total acquisition time (TT) = 36 min 12 sec, and the frame rate = 0.092 frames per second (fps).

*<u>CMR acquisition protocol for ECV</u>*: Pre-contrast T1 mapping with VFA for the heart (Fig.2B red block 1) and jugular vein (Fig. 2B blue block 1) was acquired before Gd bolus at time 0 (Fig.2B yellow block), followed by cine MRI (Fig. 2B green block) for multi-slice short-axis (SA), long-axis 2-chamber (LA2C), and long-axis 4-chamber (LA4C) imaging. Once the single steady state is reached at around ∼20 minutes post-Gd bolus, post-contrast T1 mapping for the heart (Fig. 2B red block 2) was acquired with VFA flanked by 2 jugular vein T1 mapping (Fig. 2B blue blocks 2 and 3) immediately before and after heart T1 mapping (Fig. 2B red block 2).

*<u>Quantitative T1 mapping for the jugular vein (JV) blood</u>*: Fast free-breathing-no-gating T1 mapping for a jugular vein with 4 flip angles was acquired with following parameters: FOV = 4.5 cm × 4.5 cm, matrix = 128 × 128, in-plane resolution = 0.352 mm, SLTH = 2 mm, FA = 3°, 19°, 22°, 28°, TE = 2.349 msec, TR = 77.2 msec, NR = 10, TT = 49 sec. The imaging plane was aligned with JV and completely included JV to avoid through-plane flow. The entire JV blood flow was in-slice with no through-slice flow.

*Quantitative T1 mapping for the heart*: Fast free-breathing-no-gating T1 mapping with 4 flip angles was acquired for the heart with following parameters: FOV = 5 cm × 5 cm, matrix = 256 × 256, in-plane resolution = 0.195 mm, SLTH = 1.5 mm, FA = 3°, 19°, 22°, 28°, TE = 3.06 msec, TR = 200 msec, NR = 10, TT = 2 min 26 sec.

*Cine CMR*: Fast free-breathing-no-gating cine CMR for multi-slice short-axis (SA), long-axis 2-chamber (LA2C), and long-axis 4-chamber (LA4C) cine CMR was acquired with the following parameters: FOV = 5 cm × 5 cm, SLTH = 1.5 mm for SA and 2 mm for LA, matrix = 256 × 256, in-plane resolution = 0.195 mm, TE = 3.06 msec, TR = 5.65 msec, FA = 10°, NR = 250, cardiac phases = 20, TT = 3 min 2 sec each.

### 2.6 Systolic cardiac functions and strain Analysis

Systolic cardiac functions were analyzed using the Circle cvi42 (Circle Cardiovascular Imaging, Inc.) software, such as LV volume, ejection fraction (EF), and stroke volume (SV). Strains are values that quantify the extent of ventricular deformation throughout cardiac phases: stretching/elongation or compression/shortening. Three classes of orthogonal principal strains were analyzed using Circle cvi42: circumferential, radial, and longitudinal strain for each cardiac phase.

### 2.7 Cardiac phase registration for T1 and ECV quantification

To quantify T1 at the end-diastolic (ED) phase, Singular Value Decomposition (SVD) was employed to automatically detect the end diastole (ED) phase from the Intragate VFA time series. Each VFA time series was processed independently. First, the time series was reshaped as row vectors. Then an average phase series was reconstructed by averaging the re-construction of just the first eigen value of the left and right eigenvectors. The sign of this signal was taken and the max value of the sign-time series was defined as ED.

### 2.8 T1 Mapping, ΔR1 calculation, and ECV Mapping

Increase Gd uptake resulted in a brighter MR signal in T1-weighted images. The differences in overall T1 reduction are proportional to the amount of Gd taken up by the tissue. The longitudinal relaxation rate (R1 = 1/T1), the inverse of the longitudinal relaxation time (T1), was quantitatively mapped from the IntraGate VFA images using a variable projection approach combined with non-linear least squares optimization. We first computed the R1-depenendet signal decay for the pre- and post-Gd administration inversion times. We model the MR signal’s nonlinear dependence on R1 by via a combinatory equation that combines FA effects and R1 decay into the following equation:

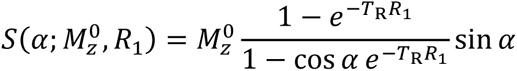

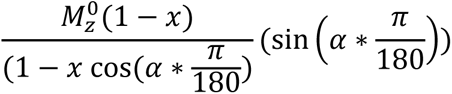

Where 𝛼 represents the FA of the image and 𝑀_z_^0^ represents the initial magnetization amplitude. The variable projection method was used to separate the linear (𝑀_z_^0^amplitude) and nonlinear (R1) parameters to reduce dimensionality during nonlinear fitting. The final R1 values are re- shaped into an image with pre-Gd and post-Gd R1.

ECV quantification was performed using the following equation:

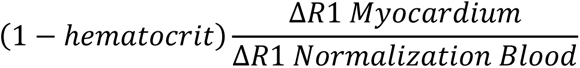

Δ𝑅1 𝑁𝑜𝑟𝑚𝑎𝑙𝑖𝑧𝑎𝑡𝑖𝑜𝑛 𝐵𝑙𝑜𝑜𝑑

For repeatability experiments, each rat underwent duplicate imaging protocols. Hematocrit is taken once for each rat and used to calculate ECV for both scans. The

Δ𝑅1 𝑁𝑜𝑟𝑚𝑎𝑙𝑖𝑧𝑎𝑡𝑖𝑜𝑛 𝐵𝑙𝑜𝑜𝑑 term is blood sourced from either the LV blood pool or jugular vein blood.

### 2.9 Ischemic reperfusion injury

Anesthesia was induced using 2.5% inhaled isoflurane in 2-month-old male rats weighing 275 ± 25 g. The rat was put on RoVent® Jr. small animal ventilator (Kent Scientific Corporation, Torrington, CT, USA). The chest of the rat was shaved. Thoracotomy was performed; the thoracic cavity was opened between 4th and 5th rib of the left side of rat. A surgical retractor was placed between the ribs to keep the cavity open. Pericardium was cleared to make the heart accessible for left anterior descending (LAD) coronary artery ligation. By gently pushing the right-side ribs, heart was temporarily brought out of the chest near the cavity opening, LAD coronary artery was located and ligated using a 6-0 surgical suture to create ischemia. A flexible rubber tubing was placed between the suture knots to safely enable the untying of knot after designated ligation time without injuring the myocardium. The left ventricle turned pale distal to ligation site immediately after inducing ischemia. After LAD ligation, heart was slid back into its position and the cavity was temporarily closed. After 60min, the ligation was released and the thoracic cavity was closed, the rats were recovered from anesthesia and housed in individual cages for 28 days to allow for the development of myocardial fibrosis.

### 2.10 Statistical analysis

Descriptive statistics was used for normally distributed variables. A combination of the D’Agnostio & Pearson and the Anderson-Darling test were used to assess normality of datasets. If the dataset failed normality, the median and interquartile range (IQR) was utilized to describe the distribution.Two-tailed paired t-test was used with α=0.05. All statistical analysis was performed in Prism (v10.2.3, GraphPad Boston, MA USA).

## 3 Results

### 3.1 Free-breathing gating-free cine and dynamic contrast enhancement (DCE) time series with a single bolus subcutaneous (subQ) Gd administration

A single bolus of Gd can be successfully administered to an anesthetized rat inside the MRI scanner via a sterile subcutaneous (subQ) catheter (Fig.1A). This enables pre- and post-contrast acquisition without moving the animal. As subQ catheters are non-invasive, this allows repetitive imaging on the same animals. Fast free-breathing gating-free cine (Fig.1B) and dynamic contrast enhancement (DCE, Fig.1C) time series can be acquired with IntraGate which convolves cardiac and respiratory motions for image reconstruction. Fig. 1B shows 8 out of the 20 cardiac phases for a mid-SA slice at the pupillary muscle level. A DCE time course (Fig. 1C) can be successfully reconstructed into a single cardiac phase (ED, end diastole) allowing visualization of the contrast-specific dynamics over time without motion artifacts.

### 3.2 Acquisition schema and DCE time courses

Dynamic contrast enhancement (DCE) signal time course of LV myocardium (Fig.2A) after a single subQ Gd bolus reached a steady state at around ∼20 min. Fast T1 mapping can be successfully acquired with IntraGate without ECG nor respiratory gating with 4 FAs (3°, 19°, 22°, 28°) for jugular vein blood (49 sec, Fig.2B blue blocks) and heart (2 min 26 sec, Fig.2B red blocks) before and after Gd bolus (Fig.2B, yellow block). It is important to acquire post-Gd contrast T1 mapping ∼20 minutes after the Gd bolus during the steady state. To prevent potential quasi-steady-state fluctuation, the post-contrast jugular vein blood T1 mapping was repeated twice, once before (Fig.2B blue block 2) and once after (Fig.2B blue block 3) the heart T1 mapping (Fig.2B red block 2). The two post-contrast jugular vein blood T1 were averaged to calculate myocardial ECV. During the waiting period for the heart signals to reach the steady state after the Gd bolus, the full cine CMR (Fig.2B green block) with 20 cardiac phases was acquired for multi-slice short-axis (SA), long-axis 2-chamber (LA2C), and long-axis 4-chamber (LA4C). The cine CMR was acquired for global systolic function and strain analysis, including LV volumes, ejection fraction, stroke volumes, and 3 orthogonal principal strains (circumferential, longitudinal, and radial).

The mean DCE time course of LV myocardium (Fig.2C) and LV blood pool (Fig.2D) across all eight rats demonstrated similar dynamics with a steady increase until 20 minutes when both reached the steady state. The overall variance across the time courses for the LV blood pool (STD LV blood pool 0.11) is higher than that of the LV myocardium (STD LV myocardium 0.078). The mean DCE time course for the jugular vein (Fig.2E) of the eight animals had a more rapid, logarithmic increase to the steady state by 15 minutes post-Gd administration and was held steady past 30 minutes. The overall variance of the JV (STD JV 0.089) is lower than that of the LV blood pool (STD LV blood pool 0.11).

### 3.3 T1 mapping with varied flip angles (VFA)

Free-breathing gating-free T1 mapping was achieved by voxel-wise T1 calculation with 4 flip angles (FA, 3°, 19°, 22° and 28°). Fig. 3 shows a mid-SA slice (Fig.3A) at the pupillary muscle level and jugular vein (Fig.3B) with different FAs before (top row) and after (bottom row) Gd contrast. We calculated the average native MR signal intensity in the LV myocardium and LV blood pool (Fig 3A) and the jugular vein (Fig 3B) for each FA. We plotted the MR intensity as a function of the flip angle (3°, 19°, 22° and 28°) and observed a logarithmic response in both the LV myocardium (Fig.3C) as well as the LV blood pool (Fig.3 D). The final 28° flip angle images showed a 1.64 and 1.28-fold increase in contrast for the LV myocardium and LV blood pool, respectively. In the jugular vein (Fig.3E) we observed a more linear increase in MR intensity as a function of the VFA with a final 1.17-fold contrast gain.

**Figure 3.**
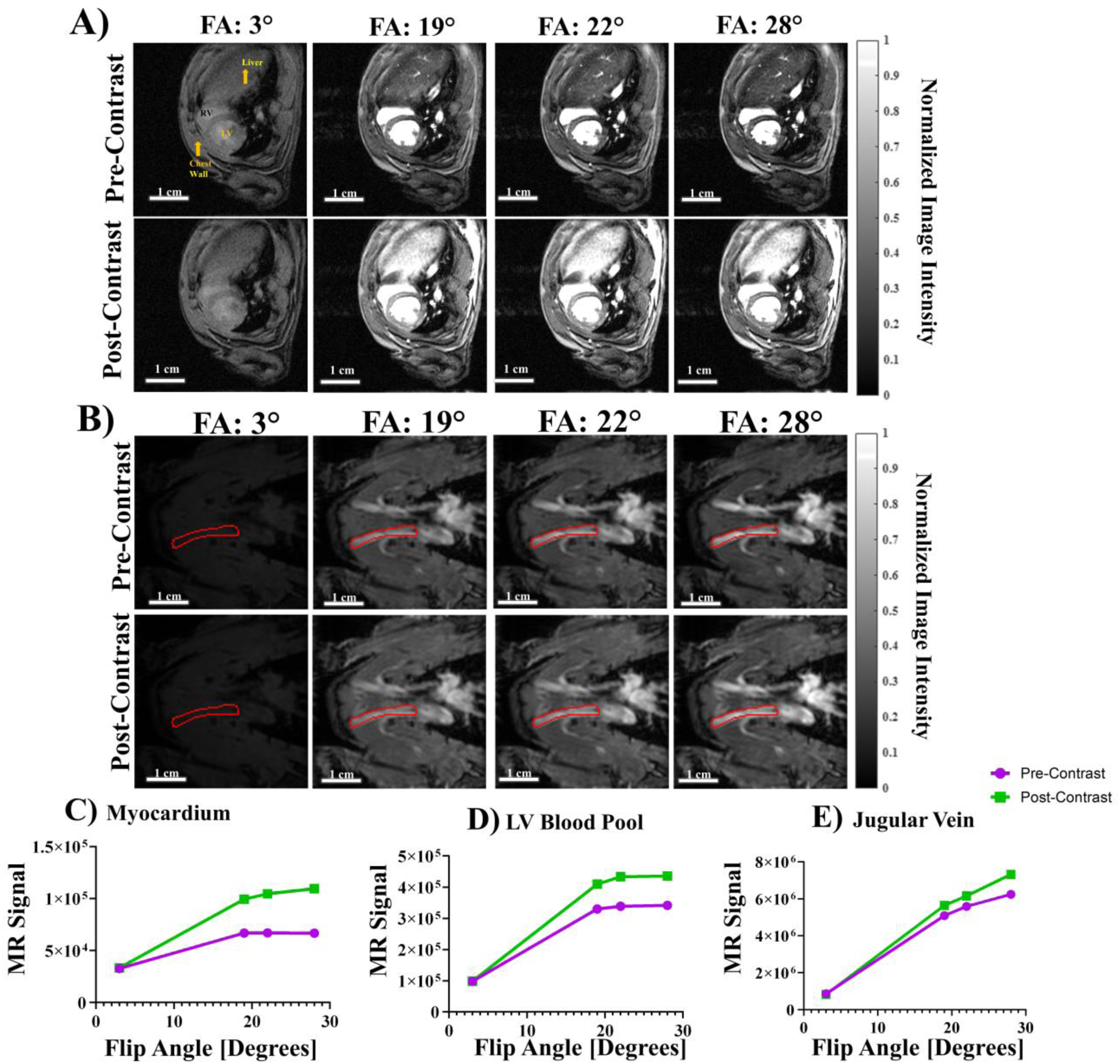
Varied Flip Angle (VFA) Imaging for cardiac structures and the jugular vein. (A) ED images for each FA for both pre- (top row) and post- (bottom row) Gd administration. All images were normalized to the max image intensity of the post-contrast FA = 28° scan to show relative changes in image intensity. (B) Jugular vein VFA images for each FA used in this study pre- (top row) and post- (bottom row) contrast. The right jugular vein is outlined in red in each image. All images were normalized to the max image intensity of the post-contrast FA 28° image to show relative changes in image intensity. The red lines outline a jugular vein. (C-E) Average signal intensity as a function of FA for both pre-Gd (purple) and post-Gd (green) series for (C) LV myocardium, (D) LV blood pool, and (E) jugular vein at 4 FA: 3°, 19°, 22, ° and 28°. We observed a consistent logarithmic response in image intensity as a function of FA.

We performed quantitative R1 mapping of the LV myocardium (Fig.4) pre- and post- Gd administration for eight animals, scanned in duplicates. Fig. 4A shows an example of LV myocardial R1 maps pre- (Fig.4A left) and post- (Fig.4A middle) Gd contrast as well as the ΔR1 map (Fig.4A right) as the differences of the two. The mean pre-contrast R1 in Sprague Dawley rat myocardial tissue was 2.31 ± 1.16 s^-1^ (Fig. 4B). The post-contrast quantitative R1 across the same scans was 3.47 ± 1.93 s^-1^ (Fig. 4C). The overall mean ΔR1 for LV myocardial tissue was 1.12 ± 0.80 s^-1^ (Fig. 4D).

**Figure 4.**
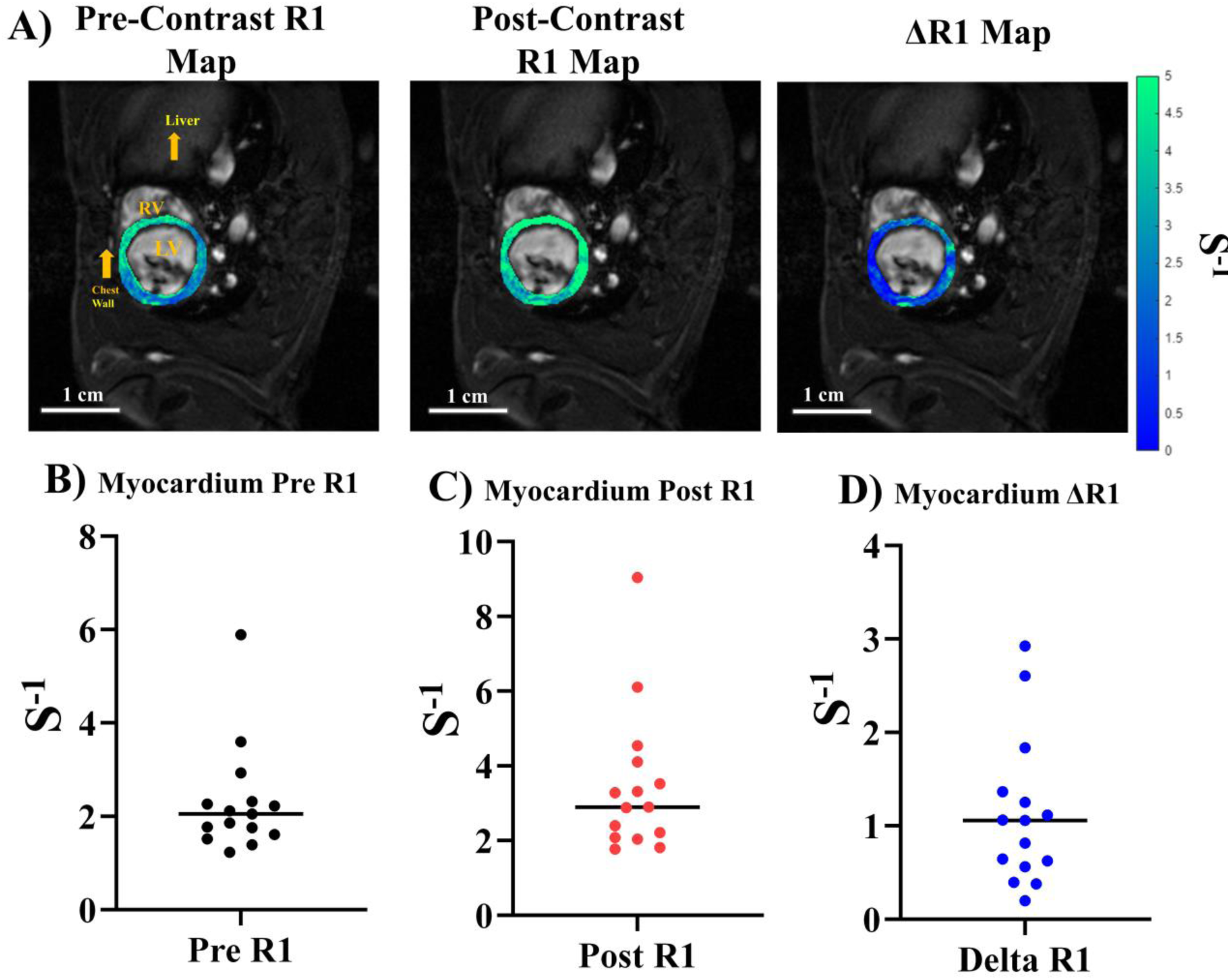
R1 mapping of LV myocardium. (A) R1 maps overlaid anatomical FA = 28° images at ED. Left: pre-contrast R1 map. Middle: post-Gd contrast R1 map. Right - ΔR1 map, the differences between the pre- and post-Gd contrast. (B-D) Median R1 plots from *n*=16 scans for (B) pre-contrast, (C) post-Gd contrast, and (D) ΔR1 of the LV myocardium for *n*=16 scans.

We performed quantitative R1 mapping of the blood pools (Fig.5) pre- and post- Gd administration for eight animals, scanned in duplicates. Fig. 5A shows an example of LV blood (top row) and jugular vein (bottom row) R1 maps pre- (Fig.5A left) and post- (Fig.5A middle) Gd contrast as well as the ΔR1 map (Fig.5A right) as the differences of the two. The LV blood (Fig.5A top row) showed higher variations in R1 and ΔR1 due to inflow turbulent and blood mixing in LV. Across the sixteen scans, we quantified the median ΔR1 for LV blood (Fig.5B) and jugular vein blood (Fig.5C). The mean ΔR1 for the LV blood pool (Fig.5B) was found to be 13.42 ± 7.15 s^-1^ whereas the mean ΔR1 for the jugular vein was 1.92 ± 0.49 s^-1^. Due to differences in the magnitude of the mean across the three datasets, the coefficient of variation was calculated to provide an un-biased measure of variability. The coefficient of variation for the LV blood pool and JV were 53.2% and 25.3% respectively. LV blood pools had much larger variations in R1 and ΔR1 due to inflow turbulent and blood mixing in the LV. Jugular vein blood had more consistent R1 and ΔR1.

**Figure 5.**
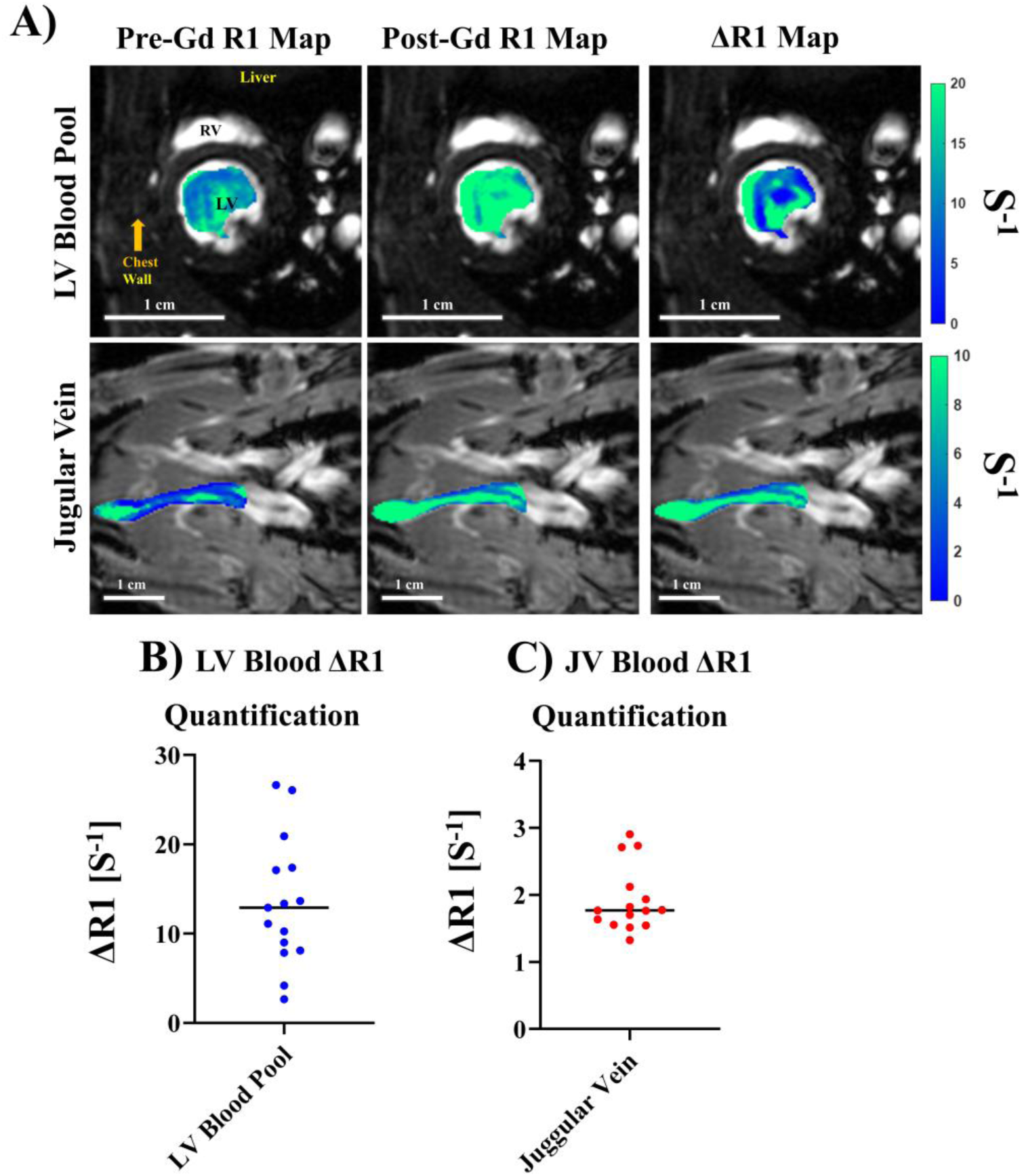
R1 mapping of blood pools. (A) R1 maps overlaid anatomical FA = 28° images at ED for the LV blood pool (top row) and the jugular vein (bottom row), showing Pre-Gd R1 maps (left column), post-Gd R1 maps (middle column) and ΔR1 maps (right column) for the left ventricle blood pool (top row), jugular vein (bottom row). (B, C) Median ΔR1 quantification for both normalization sources: (B) LV Blood Pool, (C) the Jugular Vein for *n*=16 scans.

### 3.4 ECV quantification and reproducibility

The subcutaneous catheter is non-invasive and can enable repetitive ECV quantifications.

Each rat was scanned twice on 2 different days to test the reproducibility of the ECV quantification. Myocardial ECV was calculated twice using either LV blood (Fig.6A, B) or jugular vein blood (Fig.6C, D) for blood pool normalization. Hematocrit was taken once for all eight animals, the same hematocrit value for a given animal was used to evaluate all ECV maps. The average hematocrit for the eight animals was found to be 47.4 ± 1.64 %.

**Figure 6.**
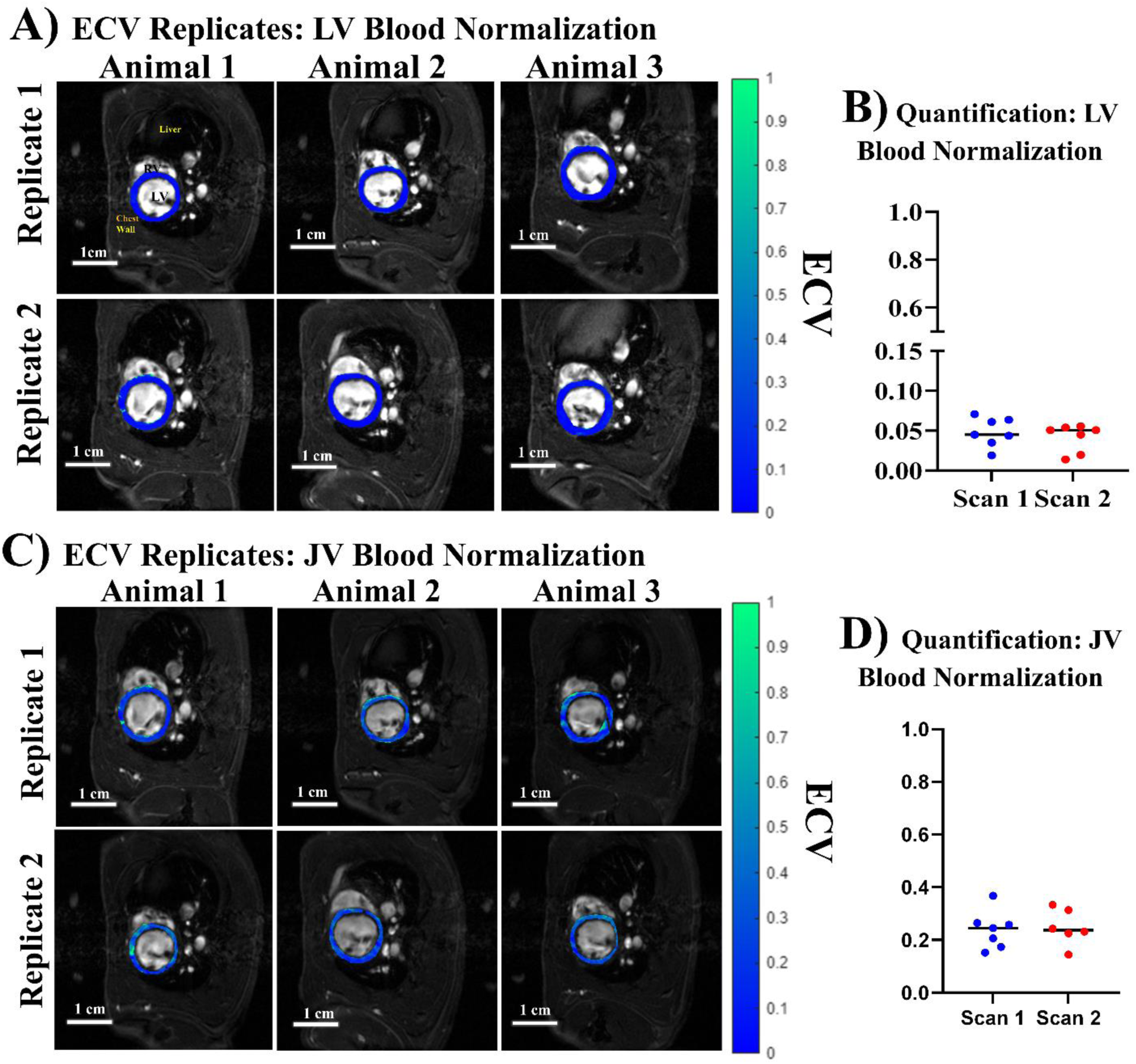
LV myocardial ECV Repeatability from different blood sources for normalization. Each rat was imaged on 2 different days to quantify ECV. ECV was calculated twice with normalization by the LV blood pool (A, B) or jugular vein blood (C, D). (A) LV myocardial ECV maps of 3 rats on day 1 (top row) or day 2 (bottom row) using LV blood pool for normalization. LV myocardial ECV maps were overlaid on anatomical FA = 28° images at ED. (B) Median ECV for LV myocardium calculated twice with normalization by the LV blood pool on 2 different days. (C) LV myocardial ECV maps of 3 rats on day 1 (top row) or day 2 (bottom row) using jugular vein blood pool for normalization. LV myocardial ECV maps were overlaid on anatomical FA = 28° images at ED. (B) Median ECV for LV myocardium calculated twice with normalization by the jugular vein blood pool on 2 different days.

First, we evaluated LV myocardial ECV using LV blood normalization (Fig.6A,B), following human myocardial ECV convention. Qualitatively, there were little differences between replicate 1 (Fig.6A top row) and replicate 2 (Fig.6A bottom row) of the same animals. We calculated the median ECV of each scan performed and evaluated differences between the means of scan 1 and scan 2 ECV distributions. We found no difference between scan 1 (Mean ECV 0.048 ± 0.018) and scan 2 (Mean ECV 0.041 ± 0.017) using paired t-testing between the two groups (*p* = 0.60). We then evaluated ECV repeatability using jugular vein blood for normalization (Fig.6 C, D). Again, qualitatively minimal differences were found from scan-to-scan (Fig. 6 D). The scan 1 and scan 2 distributions had means of 0.249 ± 0.072 % and 0.249 ± 0.068, respectively (Fig.6D). We found no difference between scan 1 and scan 2 using paired t-testing between the two groups (*p* = 0.99). The repetitive ECV quantifications in the same animals on different days showed excellent reproducibility.

The ECV quantification by LV blood and or jugular vein blood normalization differed by two main factors: magnitude of ECV and within distribution variation for each set of duplicate scans. The magnitude of ECV was ∼5 fold higher using jugular vein blood versus LV blood for normalization. The mean myocardial ECV (0.048) obtained by LV blood normalization was non-physiological, whereas mean myocardial ECV (0.25) obtained by jugular vein blood normalization was comparable to literature reports of baseline myocardial ECV in both humans [15–21] and rodents [38–42]. Additionally, the coefficient of variation across the 16 scans when utilizing LV blood pool for normalization was 39.4%, whereas when using the jugular vein blood, it was 28.1%.

### 3.5 ECV quantification for fibrosis after ischemic reperfusion injury

We tested if this ECV protocol can be applied in the pathological condition on ischemic reperfusion injury (IRI) model (Fig.7). 28 days after IRI, ECV near the IRI site was elevated (Fig.7A, yellow arrow), indicating increased myocardial fibrosis. This was validated by histological trichrome staining (Fig.7B, yellow arrow) showing blue collagen deposits in the corresponding region of the myocardium. Strain analysis (Fig.7C-F) showed compromised strain in the corresponding region of the myocardium. Our data showed that this ECV protocol can be used to detect pathological conditions.

**Figure 7.**
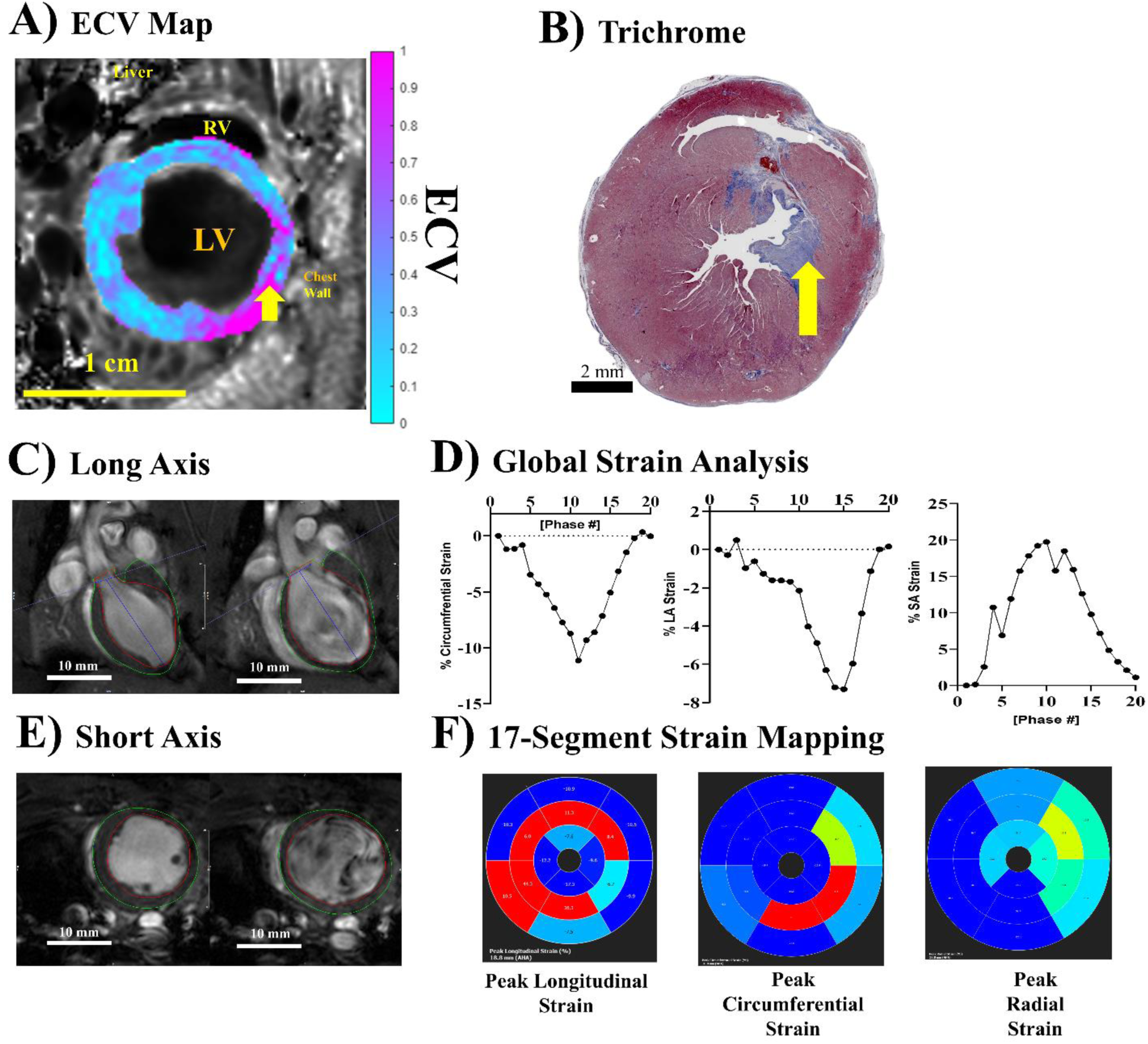
ECV mapping with ischemic reperfusion injury (IRI). (A) LV myocardial ECV map overlaid on anatomical FA = 28° images at ED using jugular vein blood pool for normalization on day 28 after IRI. The yellow arrow points to the site with high ECV reflecting high fibrosis. (B) Trichrome histology of the same animal, taken at 40x magnification and shown at 0.8x magnification. The yellow arrow points to the site with fibrotic deposition (blue), corresponding to high ECV regions in A. (C) Long Axis cine showing ES (left) and ED (right). (D) Short axis cine showing ES (left) and ED (right). (E) Global strain plots showing the % change in cardiac strain throughout the cardiac cycle for longitudinal strain (left), circumferential strain (middle plot), and radial strain (left plot). (F) 17-segment maps showing the spatial strain distribution throughout the different regions of the rat heart.

## 4 Discussion

### 4.1 Non-invasive subcutaneous protocol for ECV quantification

We have successfully established a non-invasive fast ECV quantification protocol to enable longitudinal repetitive ECV quantifications in rodents. Our protocol uses a subcutaneous catheter to deliver a Gd bolus inside the MRI scanner. Our data demonstrated that subcutaneous Gd bolus administration induced reasonable DCE time course reaching a steady state for stable T1 (or R1) quantification. Furthermore, our protocol uses free-breathing gating-free variable flip angle (VFA) T1 (or R1) quantification scheme leveraging the IntraGate sequence with retrospective gating. This allows for rapid acquisition of cardiac (∼2.5 min) and jugular vein (49 sec) T1 (or R1) quantification with no cardiac nor respiratory motion artifacts. In addition, full cine CMR can be acquired during post-Gd waiting period using IntraGate for full functional analysis. The entire full protocol for pre- and post-contrast T1 (or R1) quantification with full cine CMR can be completed within 30 minutes without ECG or respiratory gating in anesthetized rodents. This protocol is simple, robust, and highly reproducible. Our data demonstrated excellent reproducibility between 2 repeated scans on the same animals performed on different days. The non-invasive subcutaneous catheter protocol can be repeated many times. This simple non-invasive protocol can facilitate longitudinal monitoring of ECV with full cardiac function for disease progression. It can be used to monitor therapeutic interventions for disease regression in rodent models.

### 4.2 Jugular vein vs LV Blood Pool Normalization

Conventional ECV protocols [15–22] use inversion recovery schemes for T1 (or R1) quantification [32–34], such as modified Look-Locker inversion recovery (MOLLI), shortened MOLLI (ShMOLLI), and saturation recovery single-shot acquisition (SASHA). The inversion pulses remove proton spins on the acquisition planes, including the incoming blood to LV. There are usually little residual fresh spins on the acquisition planes. With the VFA method, the fresh proton spins in the incoming blood can introduce variable blood signal intensities due to inflow turbulent and fresh blood mixing in the LV. As a result, ECV normalization using the LV blood pool can be variable and inaccurate. In addition, fresh inflow proton spins can make blood T1 appear to be different, resulting in abnormally low ECV values (0.048). One way to overcome this inflow artifact is to implement a crusher gradient or a long low-power adiabatic pulse to saturate the incoming fresh spins before T1 (or R1) acquisition. However, additional crusher gradient or adiabatic pulse can prolong scan time, which can defeat the purpose of fast acquisition. In addition, inflows for the heart can come from variable directions from the pulmonary vein, superior vena, and inferior vena cava. The inflows from left atrium to left ventricle through the mitral valve can be turbulent. These can make complete saturation of incoming spins in inflows variable.

Therefore, we implemented an alternative jugular vein blood pool for normalization. The JV imaging plane was aligned with JV blood flow and included the entire JV completely, thus JV blood flow was all in-slice without through-plane flow. To avoid potential quasi-steady-state fluctuation of Gd in the blood, we acquired 2 post-contrast jugular vein T1 (or R1) quantifications immediately flanking the heart T1 (or R1) acquisition, then used the averaged of the two for ECV quantification. Our data demonstrated that LV myocardial ECV quantification using the triple jugular vein T1 (or R1) acquisitions (1 pre-contrast and 2 post-contrast) are highly reproducible between repeated scans on the same animals. Furthermore, the mean baseline ECV value (0.249) obtained in the Sprague Dawley rats using the triple jugular vein blood T1 (or R1) normalization was physiological and comparable to the reported baseline ECV values in both humans [15–21] and rodents [38–42].

## 5 Limitations

This fast non-invasive ECV protocol requires the steady state of Gd in the blood and myocardium. This protocol cannot work if the steady state cannot be achieved when the Gd dosage is too low that the subcutaneous Gd absorption is too slow and becomes the rate limiting step. Thus, optimal Gd dosage is needed for this protocol.

Using jugular vein blood for ECV quantification can mitigate the inflow artifacts due to incoming fresh proton spins draining from the great veins to the heart. However, this cannot eliminate the fresh proton spins partition into myocardium via coronary arteries. Thus, this protocol will slightly over-estimate the myocardial T1.

In order to measure blood T1 (or R1) in jugular veins as closely as possible to the time of myocardial T1 (or R1) acquisition, we used a very short jugular vein VTR protocol with only 49 seconds for all 4 FAs. This was accomplished by lower spatial resolutions. The drawback of this approach is potential partial volume artifact of the jugular veins. To prevent the partial volume artifact of jugular veins, the voxel sizes of the jugular vein T1 (or R1) acquisitions need to be optimized based on the size of the jugular veins, in particular for mice or juvenile animals.

## 6 Conclusion

We have established a simple, non-invasive, fast, and robust CMR protocol in rodents that can enable longitudinal repetitive ECV quantifications for cardiovascular disease progression and regression with interventions.

## Author contributions

**Devin R. E. Cortes:** Investigation, Methodology, Formal analysis, Software, Validation, Data curation, Writing – original draft; Writing – review & editing. **Thomas Becker-Szurszewski**: Data curation, Methodology, Formal analysis. **Sean Hartwick**: Data curation, Methodology, Formal analysis. **Muhammad Wahab Amjad**: Data curation, Writing – review & editing. **Soheb Anwar Mohammed:** Data curation, Writing – review & editing. **Xucai Chen:** Data curation, Writing – review & editing. **John J. Pacella:** Data curation, Supervision, Writing – review & editing. **Anthony G. Christodoulou:** Investigation, Methodology, Writing – review & editing. **Yijen L. Wu:** Conceptualization, Investigation, Methodology, Formal analysis, Software, Validation, Data curation, Project administration, Funding acquisition, Supervision, Writing – original draft, Writing – review & editing.

## Availability of data and materials

The datasets are available from the corresponding author upon reasonable request.

## Acknowledgements

This project was partially supported by American Heart Association (18CDA34140024), National Institutes of Health (R21EB023507), Department of Defense (W81XWH1810070), and the funds from the Rangos Research Center at UPMC Children’s Hospital of Pittsburgh.

## Abbreviation List

ECV: Extracellular Volume
CMR: Cardiac Magnetic Resonance Imaging
Gd: Gadolinium
VFA: Varied Flip Angle
DCE: Dynamic Contrast Enhancement
HCM: Hypertrophic Cardiomyopathy
CHD: Congenital Heart Disease
HFpEF: Heart Failure with Preserved Ejection Fraction
IR: Inversion Recovery
MOLLI: Modified Look-Locker Inversion Recovery
ShMOLLI: Shortened Modified Look-Locker Inversion Recovery
SASHA: Saturation Recovery Single-Shot Acquisition
ECG: Electrocardiogram
LV: Left Ventricle
JV: Jugular Vein
subQ: Subcutaneous
FLASH: Fast Low-Angle Shot
SA: Short Axis
LA2C: Long Axis 2-Chamber
LA4C: Long Axis 4-Chamber
EF: Ejection Fraction
SV: Stroke Volume
ED: End Diastolic
SVD: Singular Value Decomposition
LAD: Left Anterior Descending Coronary Artery
IRI: Ischemic Reperfusion Injury

